# Phytochemicals and Antioxidant Activity of Leaf Extract and Callus Cultures of *Cinnamomum camphora* L

**DOI:** 10.1101/2025.03.04.641517

**Authors:** Sajal Rasool, Sheza Ayaz Khilji, Zahoor Ahmad Sajid

**Author notes:** Corresponding author; Phone; 0923347407566.

## Abstract

*Cinnamomum camphora* L. is highly significant landscape tree known for its medicinal values and presence of secondary metabolites that have antioxidant, antimicrobial, anticancer, anti-inflammatory effects and widely utilized in pharmaceutical and cosmetic industry. The present work was aimed at comparative analysis of phytochemicals (phenolic and flavonoid) and antioxidant activities of calli and leaf extract of field grown camphor plant. To get *in vitro* germplasm, callus formation and direct shoot initiation was carried out and it was observed that MS medium supplemented with 0.4 µM thidiazuron (TDZ) + 2.26 µM 2, 4-dichlorophenoxyacetic acid (2, 4-D) proved best for shoot initiation from nodal explant. MS medium fortified with various plant growth regulators was used for callus formation and best callus induction response (100%) from nodal and leaf explants was observed on 2.26 µM 2, 4-dichlorophenoxyacetic acid (2, 4-D) + 8.87 µM 6-Benzylaminopurine (BAP). Callus was successfully sub-cultured and this *in vitro* proliferated callus extract and fresh leaf extract of field grown plant were used for comparative study of phytochemicals. The antioxidant enzyme activity of peroxidase (31.12 UmL^-1^ of enzyme), superoxide dismutase (35.24 Umg^-1^ of protein), and catalase (58.6 UmL^-1^ of enzyme) was higher in callus as compared to the leaf extract while glutathione peroxidase activity (0.552 Umg^-1^) was comparatively higher in leaf extract rather than callus cultures. Higher phenolic contents (1.106 mg GAE g^-1^ of FW) were recorded in callus cultures, however flavonoid contents (7.87 mg CatE g^-1^ of FW) were higher in leaf extract. This investigation showed that *in vitro* conditions and the use of plant growth regulators in various combinations might be work as elicitors to enhance the phytochemicals and antioxidant enzymes in callus culture as compared to the leaf extract.

## INTRODUCTION

Phytochemicals are the most significant group of secondary metabolites that are produced by plants for their reproduction, symbiotic relationships, growth and development (Forni *et al*., 2019). Although they are already present but stressful growth environments or modifications to the growth medium may further trigger their synthesis like conditions for *in vitro* plant (Forni *et al*., 2016; Lucioli *et al*., 2017). The majority of these chemicals are produced constitutively but their synthesis can be increased under stress in a way that it depends on the growth circumstances and the stressor (Mahajan and Tuteja, 2005; Ramakrishna and Ravishankar, 2011). Phenolic content and antioxidant activity in plant products have been shown to be positively correlated with each other (Aryal *et al*., 2019). One of the major subgroups of phytochemicals with potential antioxidant benefits and positive impacts on human health is the polyphenol (Cory *et al*., 2018) that have significant role in determining the flavour, texture, color, and sensory perception of food (Shahidi and Ambigaipalan, 2015). Phenolics are the most promising phytochemical for further studies that have vital role in the detoxification of H_2_O_2_ in plants, as well as in UV protection and enzyme modulation (Chowdhary *et al*., 2022). Some researches claim that the anticancer properties of phenolic compounds are associated to apoptosis and antioxidant properties (Nandi *et al*., 2007) and the use, identification with standardization of phytochemicals play crucial role for effective treatments (Vignesh *et al*., 2022).

Similarly, flavonoids are actively present polyphenolic phytochemicals secreted by plants (Donadio *et al*., 2021) that have been utilized in a variety of herbal remedies since ancient times as they have antiviral, anti-bacterial, anti-inflammatory, anticarcinogenic and antimutagenic properties (Roy *et al*., 2022). Phenylpropanoid pathway synthesizes flavonoid (Tajammal *et al*., 2021), while they are synthesized in response of any microbial infection or disease and mainly accumulated in the edible sections of the plants. There are many studies in literature that have been conducted on various medicinal plants to explore phytochemicals and their pharmaceutical effects on diseases. Traditional medicines made from a variety of medicinal plants are widely used in countries; Pakistan, China, India, Bangladesh, Korea, Taiwan, Japan, and Sri Lanka. Among many of medicinal plants *C. camphora* bearing with high significance because of its herbal and ritual use especially in China and Sub-continent. *Cinnamomum camphora* L. Presl is a perennial tree that belongs to the family *Lauraceae*. It is naturally cultivated as a landscaping tree, and is used as an important herbal medicine in southern China (Tian *et al*., 2020; Zhang *et al*., 2020). It is also known as Camphor tree, Camphor, Camphorwood or Camphor laurel, and found in temperate to subtropical regions of East Asian countries i.e., China, Japan, Korea, Vietnam, especially along the coast from Cochin China (Vietnam) and also belongs to the estuary of the Yangtzekiang (river), adjoining islands; Hainan and Taiwan and the naturally grown range bounds to just about 10-36°N and 105-130°E (CABI, 2022). *C. camphora* is a well-known tree species of great significance to human beings. Leaf essential oil (CEO) has potent antioxidant, antimicrobial, anticancer, strong insecticidal (Luo *et al*., 2021) and anti-inflammatory effects (Chen *et al*., 2020). The CEOs are being widely utilized in pharmaceutical, food and cosmetics industries as significant raw resources (Ni *et al*., 2021). Camphor is helpful in evaluating more possible pharmaceuticals and creating new anti-inflammatory medications (Muhammad *et al*., 2019). In Chinese tradition, camphor leaves may treat gastrointestinal problems like diarrhea as well as mental problems like hysteria, anxiety, and neuralgia (Lee *et al*., 2006) and species of the genus *Cinnamomum* are also utilized as significant condiments (Wang *et al*., 2020a). According to other studies, ethanolic extract from leaves of camphor plant is significantly effective in treating atopic dermatitis (Kang *et al*., 2019). It has been showed that leaves of *Camphora* have remarkable anti-inflammatory capability in adult human (Wang *et al*., 2020b). Extracted essential oils from various parts of camphor tree like twigs, leaves, and seeds, has notable insecticidal potential, camphor leaves essential oil is additionally used as fruits and vegetables preservative and is predicted to have a lot of other applications. In future, the researcher hopes to develop bio-based products that will benefit from using camphor leaves in medicine (Wei *et al*., 2022). Besides this huge pharmacological, industrial and environmental significance, Camphor is not propagated at large scale in Pakistan due to several issues regarding its plantation. Camphor is exotic species in the area of sub-continent and there’s a problem with the traditional methods of propagation for instance from seeds propagation. Seeds are produced in huge amount within fruits (approximately up to 100,000 seeds per adult tree) but in Pakistan production of flowers are rare and non-viable and hence seeds are not produced. Cuttings and layering are also methods that are frequently used in propagation but the limitations of resources and poor response prevent these techniques from being widely utilized at commercial level. Consequently, the conventional methods of breeding which are described above are no longer be able to satisfy the rising demand of *C. camphora* seedlings in therapeutical and pharmaceutical industries. While on the other hand, because of its versatile therapeutic properties, the annual demand of camphor is rising day by day. The global camphor market predicts that between 2020 and 2025 its demand will expand at compound annual growth rate of 11-13% (https://www.lucintel.com/camphor-market.aspx).

Plant tissue culture has been widely used for the multiplication of this important tree species because this technique has very high regenerative efficiency. Moreover, in micropropagation, there are no limitations of climatic conditions, seasonal affect and regional restrictions, although application of this technique is very limited in regard to camphor tree. *In vitro* propagation by plant tissue culturing of camphor is challenging because of its slow growth rate. A desirable method for plant regeneration and for conservation of genetic diversity as well as use of genetic resources like bioactive substances is tissue culturing technique (Bhojwani and Dantu, 2013). For this purpose, the growth of plant is regulated by utilizing PGRs (plant growth regulators) in various concentrations and combinations while using various explants (nodal, leaves, internodal). Externally applied PGRs have potential to alter the physiology as well as internal polarity of explant. *C. camphora* can be regenerated by two pathways; somatic embryogenesis or organogenesis under controlled conditions. Based on the mentioned methods and strategies, there is a dire need to consider *in vitro* propagation strategies for this economically significant medicinal plant. Environmental conditions also effects on the phytochemical compositions of plants as well as the antioxidant capacity (Samaniego *et al*., 2020). However, the composition of phytochemicals may vary with plant species based on the specific effect and type of plant growth regulators (Lee *et al*., 2020).

Because of above mentioned *Camphora’*s medicinal value with rich source of phytochemicals and high antioxidant properties it provides an ideal system for biotechnological research. Callus cultures have significant importance for studies and understanding of secondary metabolites. So, the main objective of current research was to establish callus cultures for examining the comparative changes in phytochemicals and antioxidant activity of these calli and fresh leaf extract of camphor.

## MATERIALS AND METHODS

### Plant material and culture conditions

Young nodes and fresh juvenile leaves of *Cinnamomum camphora* L. were used as explants during this research work. The nodal segments were procured during November and December, 2022 from a healthy 20-year-old tree of Camphor grown at University of Punjab, Lahore. Murashige and Skoog (MS) medium (1962) and Woody Plant Medium WPM (1981) with 30 gL^-1^ sucrose, 8.0 gL^-1^ agar were used with different concentrations of 2, 4-D (2, 4-dichlorophenoxyacetic acid), BAP (6-Benzylaminopurine), NAA (1-Naphthaleneacetic acid), KIN (Kinetin), IBA (Indole butyric acid), TDZ (Thidiazuron), IAA (Indole-acetic-acid), Zeatin for this experiment. Explants were cleaned with running tap water 4 to 5 times to remove all the particles of dust. They were then dipped in a solution of household detergent for 15 minutes while being continuously stirred. After rinsing with distilled water these segments were treated with 15% (*v/v*) bleach for 15 minutes with gentle agitate and washed with double distilled water in laminar air flow chamber. Further treating these explants with 0.1% solution of mercuric acid (HgCl_2_), depending on maturity of explant, then washed with double distilled water for 4 to 5 times then immersed in 70% ethanol for 1 minutes and then blot-dried using sterilized blotting paper after rinsing again with distilled water. These explants were then placed one by one into the test tube containing medium while keeping them near spirit lamp to avoid any contamination. These cultures were then kept under 16/8-hours photoperiod (40 μmol m^-2^ s^-1^) and 8 hours dark period provided by white light (36Watt) at 25 ± 2°C temperature in the culture room.

### Data collection for morphological attributes

Data were recorded in terms of callus texture, days and frequency of calli formation, growth index and color of calli and also for the number of shoots, percentage response and number of leaves for all the tested media.

### Shoot induction and root formation in MS and WPM medium

Nodal segments were used as an explant in MS and WPM supplemented with various combinations and concentrations of plant growth regulators for shoot induction and root formation. The detail of various combinations used as follows: M0 (MS medium without PGRs); M1 (2, 4-D 2.26 µM + BAP 8.87 µM); M2 (2,4-D 4.52 µM + KIN 4.65 µM) ; M3 (BAP 4.44 µM + KIN 4.65 µM); M4 (TDZ 0.4 µM + 2,4-D 2.26 µM); M5 (Zeatin 4.56 µM + KIN 4.65 µM); M6 (IBA 4.92 µM + AC 2.0 g); M7 (IAA 2.46 µM + BAP 8.87 µM) and same concentrations and combinations of PGRs were used for WPM. After inoculation the data were recorded for % response (Eq. 1), number of shoots and leaves, days for shoot initiation and leaves formation.

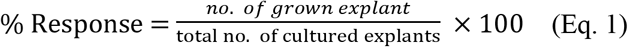

### Callus initiation and its proliferation

Leaves and nodal segments were used as explants for initiation of callus in MS media supplemented with various concentrations of plant growth regulators, C0 (MS medium without PGRs); C1 (2,4-D 2.26 µM + BAP 8.87 µM); C2 (Zeatin 4.56 µM + KIN 4.65 µM); C3 (BAP 13.3 µM +KIN 4.65 µM); C4 (TDZ 0.4 µM + BAP 8.87 µM L^-1^); C5 (IBA 2.46 µM +NAA 2.69 µM) used for callus induction from leaf while M0 (MS medium without PGRs); M1 (2, 4-D 2.26 µM + BAP 8.87 µM); M2 (2,4-D 4.52 µM + KIN 4.65 µM); M3 (BAP 4.44 µM + KIN 4.65 µM); M4 (Zeatin 4.56 µM + KIN 4.65 µM); M5 (IBA 2.46 µM + BAP 4.44 µM). After 30 days of inoculation the data were recorded for % frequency of callus formation (Eq 2), Growth index (Eq. 3), days for callus initiation, callus texture and color. Initiated calli were sub-cultured on these three media, S1 (2, 4-D 2.26 µM + BAP 8.87 µM); S2 (2, 4-D 2.26 µM + TDZ 0.4 µM); S3 (IAA 9.84 µM + BAP 8.87 µM) with the interval of 20 days to maintain healthy calli.

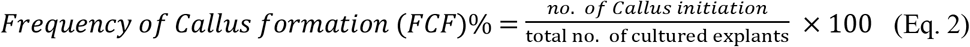

The fresh weight of callus formation/growth index was measured using following formula:

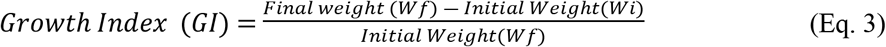

*Wi*= initial weight.

*Wf*= final weight.

### Comparative Analysis of Antioxidant Enzyme Activities and Phytochemicals of Leaf Extract and Callus Culture of *C. camphora* L

#### Extraction and estimation of antioxidant enzymes

Healthy and fresh leaves of Camphor were collected and packed in polythene bags and brought to Plant Developmental and Regenerative Biology Laboratory, Institute of Botany, University of the Punjab, Lahore. The plant material was rinsed under running tap water thoroughly and dried with the help of tissue paper by gentle tapping. Fresh leaves were measured 4g and placed in -80°C for 24 hrs. 4g calli were used for the extraction of enzymes. Plant material and calli were crushed with the help of pestle mortar separately. Phosphate buffer (7.2 pH) was used with 1:2 of crude extract and 0.1g PVP (*Polyvinyl polypyrrolidone*; Sigma Aldrich) was added to make homogenous slurry. Slurry was centrifuged at 4°C and 14000 rpm for 15 minutes. After centrifugation, supernatant was used for estimation of antioxidant enzyme activities and phytochemical analysis. Extraction percentage was 50% by using following formula (Eq. 4):

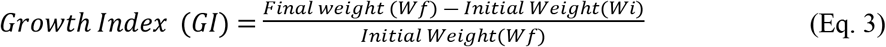

*W*1 = weight of crude extract

*W*2 = weight of sample

#### Estimation of peroxidase (E.C 1.11.1.7) activity (POD)

The ‘Guaiacol-H_2_O_2_’ method described by Luck (1974) was modified for quantitative estimation of peroxidases. Two test tubes were used, one with a reaction mixture; 3.0 mL of phosphate buffer (0.1M; pH 7.2) with 0.05 mL of guaiacol (20 mM; 2-methoxyphenol) and 0.1 mL of crude enzyme extract. Instead of crude enzyme extract, 0.1 mL of distilled water was used in the second test tube. In both test tubes, 0.03 mL H_2_O_2_ (12.3 mM) was added for the reaction in both test tubes. Peroxidase activity was estimated as the required time for increase in 0.1 (e.g., 0.4-0.5) absorbance at 240 nm and represented as UmL^-1^ of enzyme (Eq. 5).

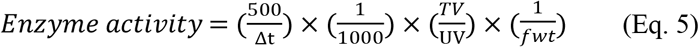

Δt = change in time; TV = total prepared volume (mL); VU = volume of the sample used (mL); fwt = fresh weight (g).

#### Estimation of catalase (E.C 1.11.1.6) activity

The activity of catalase was measured using the method described by Beers and Sizer, (1952), with some modifications. The reaction was conducted with two buffer solutions (A and B). Reagent A contained 50 mM potassium phosphate at pH 7.0 while B was 0.036% H_2_O_2_ solution with 50 mM potassium phosphate buffer. The reaction mixture of catalase was composed of 0.1 mL of enzyme extract and 2.9 mL of Reagent B, while the blank contained only 3.0 mL of Reagent A. The enzyme activity was determined by measuring the required time for decrease in absorbance from 0.45 to 0.40 (at 240 nm) and was expressed as UmL^-1^ of enzyme (Eq. 6).

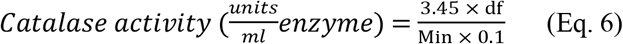

3.45= Decomposition of 3.45 micromoles of hydrogen peroxide in a 3.0 ml. of reaction mixture that produce decrease in the A_240nm_ from 0.45 to 0.40 units.

df = dilution factor (mL); Min= Time in minutes required for the A_240nm_ to decrease from 0.45 to 0.40 absorbance units; 0.1= Volume of enzyme used (mL).

#### Estimation of superoxide dismutase (E.C 1.15.1.1) Activity (SOD)

The activity of superoxide dismutase was measured by using a modified method developed by Maral *et al*. (1997). Reaction mixture was prepared by adding 1 mL of sodium cyanide (NaCN), 10 ml of Methionine, 10 mL of EDTA, 1mL of NBT (Nitroblue tetrazolium), and 1 mL of Riboflavin as a substrate. The volume of the mixture was made up to 100 mL by adding buffer solution. Two test tubes (15 × 150 mm) were used for the assay; one contained 5 µL of extract and reaction mixture, while the other tube served as a blank, contained 2 mL of reaction mixture only. Both the tubes were placed under fluorescent bulbs (30 Watt) for 15 minutes. By using spectrophotometer (UV-4000S), the absorbance of both test tubes was measured at 560 nm using a. SOD activity was further calculated on the base of this fact that 50% inhibition is caused by one unit of SOD. SOD activity was expressed as units mg^-1^ of protein. The % inhibition of NBT was calculated to determine SOD as follows (Eq. 7):

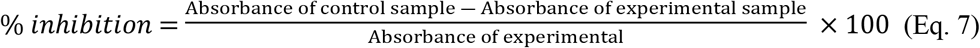

#### Estimation of glutathione peroxidase (E.C 1.11.1.9) activity (GPX)

To measure the activity of GSH-Px, the method of Flohé and Günzler, (1984) was slightly modified by using H_2_O_2_ (Duksan reagent) as a substrate. To conduct the enzyme reaction, 200 µL of the supernatant was mixed with 400 µL of 0.1mM GSH (reduced glutathione, Sigma) and 200 µL of Na_2_HPO_4_ (0.067 M) in a test tube. For the non-enzyme reaction, the same reagents were used without the supernatant extract. The mixture was pre-heated on a water bath at 25°C temperature for 5 minutes, followed by the addition of 200 µL of 1.3 mM H_2_O_2_ to initiate the reaction. This reaction was lasted for 10 minutes and then stopped by adding 1 mL of 1% trichloric acetic acid (TCA; Merck) and placing the mixture into an ice bath for 30 minutes. The supernatant was collected by centrifuging the mixture for 10 minutes at 3000 rpm, 480 µL of the supernatant was added into a test tube with 2.2 mL of Na_2_HPO_4_ (0.32 M) and 0.32 mL of 1.0 mM 5,5-dithio-bis (2-nitrobenzoic acid; DTNB) was added for development of color. The absorbance was measured at 412 nm with a spectrophotometer within 5 minutes. GPX activity is expressed as Umg^-1^. Residual glutathione concentration could be calculated by using standard curve of the glutathione. GPX activity is equals to the no. of µmol consumed glutathione (Eq. 8).

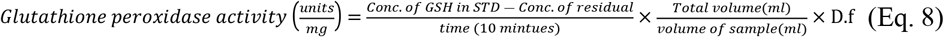

One unit of GPX is defined as the amount of enzyme that has capability to oxidize 1.0 µM GSH to GSSG per minute at temperature 25°C.

#### Estimation of total phenolic contents

Chun and Kim, (2004) method was used to determine the total phenolic content involved mixing 0.2 mL of the extract with 2.6 mL of distilled water and adding the 0.2 mL of Folin-Ciocalteu reagent after 5 minutes, then 2 ml of 7% Na_2_CO_3_ was added and stirring for 30 seconds. This solution left in the dark for almost 90 minutes and then absorbance was measured at 750 nm against blank (without enzyme extract). The amount of total phenolic contents was measured in mg GAE g^-1^. This measurement was obtained by using a standard curve Gallic acid calibration at 750 nm (Eq. 9).

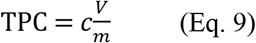

c = concentration of gallic acid obtained from calibration curve in mg/mL

v = volume of extract in mL; m = mass of extract in gram

### Determination of total flavonoid contents

To measure the amount of flavonoids, Ivanova *et al*. (2011) method was employed that involved using the AICI_3_-NaNO_2_-NaOH complex. The test tube contained 0.2 mL of the extract with 3.5 mL of distilled water, 0.15 mL of 5% NaNO_2_, 0.15 mL of 10% AlCl_3_, and 1 mL of 1 M NaOH, with the interval of 5 minutes. The reaction mixture was left to react for 15 minutes, and the absorbance was measured at 510 nm. Flavonoid content was reported in mg CatE g^−1^, which was calculated from the standard curve of catechin calibration at 510 nm (Eq. 10).

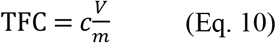

c = concentration of catechin obtained from calibration curve in mg/mL v = volume of extract in mL; m = weight of plant extract in g

### Experimental layout and statistical analysis

All experiment was performed with 10 replicates for each treatment of calli induction and shoot formation. For the analysis of phytochemical contents and antioxidant activity 3 readings were recorded for each sample and treatment. By using statistical program SPSS (Version 25.0.0), all the recorded data were analyzed to find out means and standard errors through conducting ANOVA. LSD and Duncan’s Multiple Range test was also used to find statistically significant differences among the means of various treatments. Differences were considered as significant at *P* ≤ *0*.*05*.

## RESULTS

### Standardization of the medium for shoot initiation of *c. camphora*

Young juvenile explant of camphor was collected during the months of November and December 2022. Nodal sections were inoculated on MS media containing various combinations of growth regulators. *In vitro* plants were raised to get healthy germplasm under the controlled environmental conditions. Explant treated with 10% bleach gave best surface sterilization of explant as compared to other sterilant used during this investigation and only 20% contamination was observed. For *in vitro* establishment of plant only nodal sections were used as explant during this investigation. After reviewing literature, two types of media MS and WPM having various growth regulators were tried alone and in combination with PGRs but only MS media fortified with different concentrations of PGPRs were proved effective. Shoot formation and callus induction were not observed on WPM media. So, here presented results are regarding to MS media.

### Shoot initiation of Camphor in MS medium supplemented with different plant growth regulators

For shoot initiation of camphor, MS media were used with various combinations of PGRs and activated charcoal (2.0 gL^-1^) was also used for reducing hyperhydricity (vitrification). Eight media were used for shoot initiation from nodal segment. Best results were observed on M1 medium with 100% response (Figure 1C), although it took more time (24.67 days) to initiate shooting after inoculation but it produced more number of leaves (11.00) and shoots (1.33) as compared to any other media. Similarly, M4 medium also showed 100% response within 18.00 days but number of leaves (6.33) and number of shoots (1) were less (Figure 1D) as compared to M1. Shoot initiation response was also 100% on M6 media within 19 days of inoculation. Here, the number of leaves (5.66) and number of shoots (1) were also less as compared to M1 medium (Figure 1B). M7 showed 60% response after 14.33 days of inoculation and produced 4.33 leaves and 0.67 shoots (Figure 1A). M0, M2, M3, and M5 could not support any response regarding to shooting during this investigation (Table 1).

**Table 1:** Shoot initiation from nodal segment of *C. camphora* on MS medium supplemented with various concentrations of plant growth regulators.

| Medium MS | PGRs | Concentrations (μM) | Days for Shoot Initiation | % Response | No. of Leaves | No of shoots |
| --- | --- | --- | --- | --- | --- | --- |
| M0 | 0 | 0 | 0 | 0 | 0 | 0 |
| M1 | 2, 4-D+BAP | 2.26+8.87 | 24.67 ± 1.76<sup>a</sup> | 100 | 11.00 ± 2.64<sup>a</sup> | 1.33 ± 0.33<sup>a</sup> |
| M2 | 2, 4-D+KIN | 4.52+4.65 | 0 | 0 | 0 | 0 |
| M3 | BAP+ KIN | 4.44+4.65 | 0 | 0 | 0 | 0 |
| M4 | TDZ+2, 4-D | 0.4+2.26 | 18.00 ± 1.52<sup>ab</sup> | 100 | 6.33 ± 1.45<sup>b</sup> | 1.00 ± 0.00<sup>ab</sup> |
| M5 | Zeatin+ KIN | 4.56+4.65 | 0 | 0 | 0 | 0 |
| M6 | IBA+AC | 4.92+ 2.0g | 19.00 ± 0.57<sup>ab</sup> | 100 | 5.66 ± 0.88<sup>b</sup> | 1.00 ± 0.00<sup>ab</sup> |
| M7 | IAA+BAP | 2.46+8.87 | 14.33 ± 7.17<sup>b</sup> | 60 | 4.33 ± 2.96<sup>bc</sup> | 0.67 ± 0.33<sup>b</sup> |

Values are the mean ± S.E of ten replicates for each treatment. Different alphabetical letters in each column shows the significant differences at $P \leq 0.05$ according to Duncan's multiple range test.

**Figure 1.**
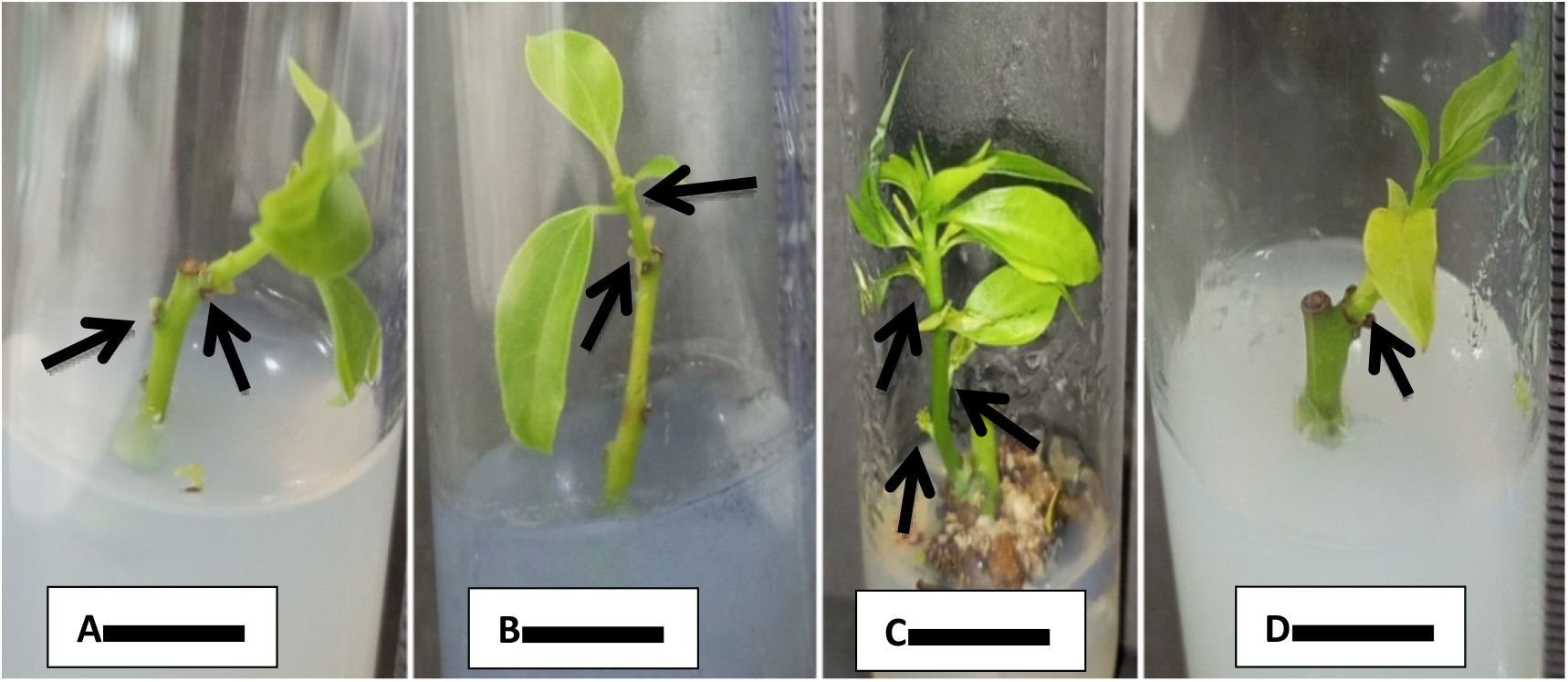
**A:** Shoot initiation from nodal segment on MS medium fortified with BAP and IAA (8.87 µM +2.46 µM) after 22 days. **B:** On MS basal media fortified with IBA (4.92µM) and AC 2.0g after 20 days. **C:** On MS basal media fortified with BAP and 2, 4-D (8.87µM+ 2.26µM) after 30 days. **D:** On MS basal media fortified with TDZ and 2, 4-D (0.4µM + 2.26µM) after 22 days. Bar = 1 cm

### Standardization of Medium for Callus Induction and Maintenance of *C. camphora*

MS media in combination with various growth regulators were tried for callus induction and proliferation of *C. camphora*. Seven media were used for callus induction from leaf explants as mentioned in Table 2. From these best media was for further utilized for proliferation of callus.

**Table 2:** Callus induction from leaf explant of *C. camphora* in MS medium fortified various combinations of plant growth regulators.

| Medium | PGRs | Concentration (μM) | Days to callus induction | FCF % | Morphology |
| --- | --- | --- | --- | --- | --- |
| C0 | 0 | 0 | 0 | 0 |  |
| C1 | 2,4-D+BAP | 2.26+8.87 | 26.2 ± 1.46<sup>b</sup> | 100 | Brownish in color with dark patches, friable and compact |
| C2 | Zeatin+ KIN | 4.56+4.65 | 0 | 0 |  |
| C3 | BAP+KIN | 13.3+4.65 | 0 | 0 |  |
| C4 | TDZ+BAP | 0.4+8.87 | 57.2 ± 1.39<sup>a</sup> | 100 | Soft callus, Whitish Yellow, friable |
| C5 | IBA+NAA | 2.46+2.69 | 0 | 0 |  |
| C6 | IAA+BAP | 9.84+8.87 | 0 | 0 |  |

Values are the mean ± S.E of ten replicates for each treatment. Different alphabetical letters in each column shows the significant differences at $P \leq 0.05$ according to Duncan's multiple range test.

### Effect of different combinations of plant growth regulators on the leaf explant for callus induction

From all the seven media only two were proved effective for callus induction with 100% response. C1 medium showed best results as compared to C4 with regards to days for callus induction. In case of C1 medium, calli were induced within 26.2 days after inoculation (Figure 2B). Morphologically, callus was brownish in color with dark patches on it, and friable in texture. Callus was induced after 57.2 days on C4 culture medium (Figure 2A). Callus produced on this medium was soft, whitish yellow in appearance and friable in texture. The entire calluses were produced from the leaf discs on both C1 and C4 mediums were non-embryogenic.

**Figure 2.**
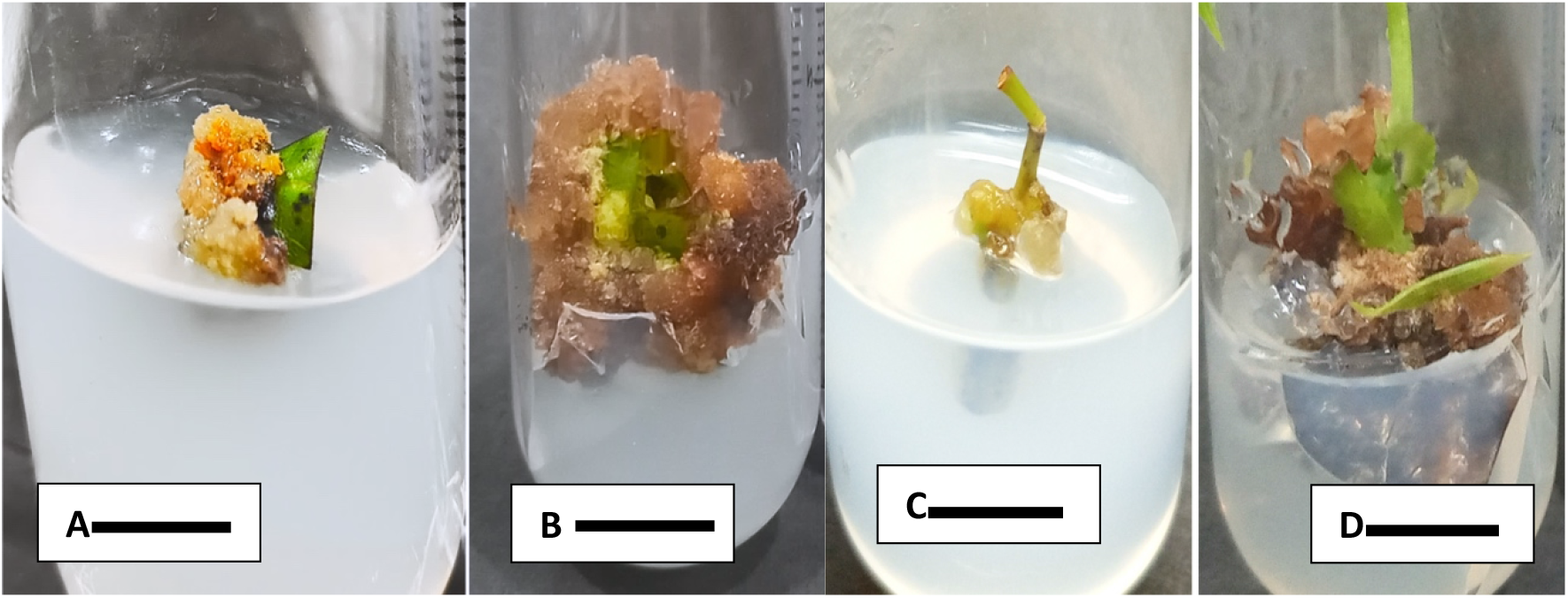
**A:** Callus induction from leaf on MS medium with TDZ and BAP (0.4µM+ 8.87µM) after 57 days. **B:** On MS medium with BAP and 2, 4-D (8.87 µM + 2.26 µM) **C**: MS medium supplemented with BAP and IBA (4.44µM+ 2.46 µM) after 22 days. **D:** Callus induction in MS medium supplemented with BAP and 2, 4-D (8.87µM+ 2.26 µM) after 30 days. Bar = 1 cm

**Figure 3.**
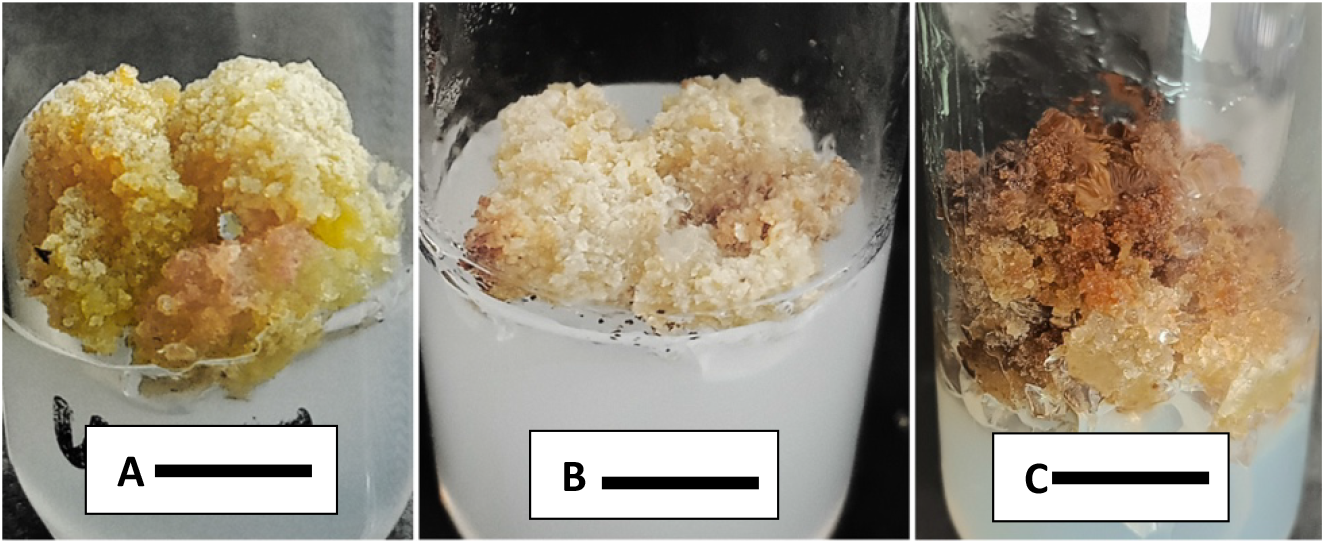
**A:** Callus proliferation in response of sub-culturing on S1 MS medium with BAP + 2, 4-D (8.87µM+ 2.26µM) after 20 days. **B:** On S3 MS medium with IAA + BAP (9.84µM + 8.87µM L^-1^) after 20 days. **C:** On S2 MS medium with TDZ+ 2,4-D (0.4 µM+ 2.26 µM) after 20 days. Bar= 1cm

### *In Vitro* callus initiation and maintenance from nodal segment and juvenile leaves of *C. camphora*

Different combinations and concentrations of PGRs were supplemented to MS (basal medium) for initiation of callus from nodal segments and young leaves of camphor plant. These cultures were incubated at temperature 25 ± 2°C under a 16-h of photoperiod, with white light intensity provided by fluorescent tube light (40 µmol^−1^ m^−2^ s^−1^). After 28 to 30 days, induced calli were further sub-cultured on the S1, S2 and S3 media. The incubation period of sub-cultured calli was 30 days but data was collected after 20 days.

### Callus induction from nodal segment of *C. camphora* in MS medium supplemented with plant growth regulators

From six media used for callus induction from nodal explants, calli was induced in two combinations (M1 and M5) of hormonal medium. M1 media showed 100% callus induction after 22.70 days of inoculation. Calli derived from M1 medium were brownish yellow and leathery in texture after 22 days of inoculation, but after maintaining, it turned into dark brown and friable texture (Figure 2D). It was observed that the hard calli produced from nodal segment in the presence of 2, 4-D and BAP were non-embryogenic with no regenerative ability.

Callus Initiation was supported about 60% in M5 media within 17.40 days but the rate of callus induction was slowest as compared to M1. Calli derived from M5 were also non-embryogenic. The deceleration in growth was observed after 50^th^ days of inoculation. However, on M5 medium calli were creamy yellowish and leathery in texture (Table 3)

**Table 3:** Callus initiation from stem segment in MS medium supplemented with plant growth regulators.

| Medium MS | PGRs | Concentration (μM) | Days to callus initiation | % Response | Callus morphology |
| --- | --- | --- | --- | --- | --- |
| M0 | 0 | 0 | 0 | 0 | 0 |
| M1 | 2, 4 D+BAP | 2.26+8.87 | 22.70 ± 0.50<sup>a</sup> | 100 | Hard callus, brownish yellow and compact in texture |
| M2 | 2,4-D+KIN | 4.52+4.65 | 0 | 0 |  |
| M3 | BAP+ KIN | 4.44+4.65 | 0 | 0 |  |
| M4 | Zeatin+ KIN | 4.56+4.65 | 0 | 0 |  |
| M5 | IBA+BAP | 2.46+4.44 | 17.40 ± 7.11<sup>b</sup> | 60 | Soft callus, creamy yellowish in color and leathery in texture |

Values are the mean ± S.E of ten replicates for each treatment. Different alphabetical letters in each column shows the significant differences at $P \leq 0.05$ according to Duncan's multiple range test.

**Table 4:** Callus Proliferation from Sub-culturing on MS medium supplemented with various combinations of Plant Growth Regulators.

| Medium | PGRs | Concentration (μM) | Days to callus growth | Growth index (g) | Morphology |
| --- | --- | --- | --- | --- | --- |
| S1 | 2,4-D+BAP | 2.26+8.87 | 19.20 ± 1.39<sup>a</sup> | 0.76 ± 0.06<sup>a</sup> | Soft callus, lush green in color,lightly friable, granular and compact in texture |
| S2 | 2,4-D+TDZ | 2.26+0.4 | 18.40 ± 1.63<sup>a</sup> | 0.47 ± 0.13<sup>a</sup> | Reddish brown with creamy base, light compact and friable |
| S3 | IAA+BAP | 9.84+8.87 | 18.40 ± 0.58<sup>a</sup> | 0.66 ± 0.10<sup>a</sup> | Whitish in color with light brown patches, granular in texture |

Values are the mean ± S.E of ten replicates for each treatment. Different alphabetical letters in each column shows the significant differences at $P \leq 0.05$ according to Duncan's multiple range test.

### Sub-culturing of callus for maintenance and proliferation

Successful sub-culturing of callus produced from nodal segment was done on three media; S1, S2 and S3. Combinations and concentrations on S1 medium were same as used before for nodal callus while two new media were S2 and S3 for further proliferation of callus cultures. S1 medium supported the production of healthy calli within 19.20 days. Morphologically, calli were soft and lush green in color, lightly friable, granular and compact in texture. Callus proliferation on S2 and S3 media took 18.40 days after inoculation. Calli in S2 media were reddish brown in color with creamy base, lightly compact and friable in texture. On S3 media calli were whitish in color with light brown patches on it and granular in texture. For achieving 0.5g of callus culture for further study, the highest growth index 0.76 g was recorded on S1 media as compared to S2 with 0.47 g and S3 with 0.66 g.

### Comparative study of antioxidant enzymes and phytochemicals of camphor leaf extract and callus cultures

#### POD, CAT, SOD and GPX activity

A Decreasing trend in the activity of peroxidases was recorded from 32.124 to 5.66 UmL^-1^ of enzyme in callus and leaf extract, respectively. Overall, POD activity of callus culture was found higher as compared to leaf extract of camphor. Similar trend was recorded in the activity of catalases which was decreased in leaf extract from 58.6 to 18.683 UmL^-1^ of enzyme. SOD activity of callus cultures was 7.078 Umg^-1^ of protein while in case of leaf extract 2.232 Umg^-1^ of protein was recorded. Unlike the above mentioned three primary antioxidants, GPX activity of leaf extract was higher as compared to callus cultures grown under the controlled conditions. Decreasing trend in the activity of GPX was from 0.552 to 0.379 Umg^-1^ of protein (Figure 5).

**Figure 4:**
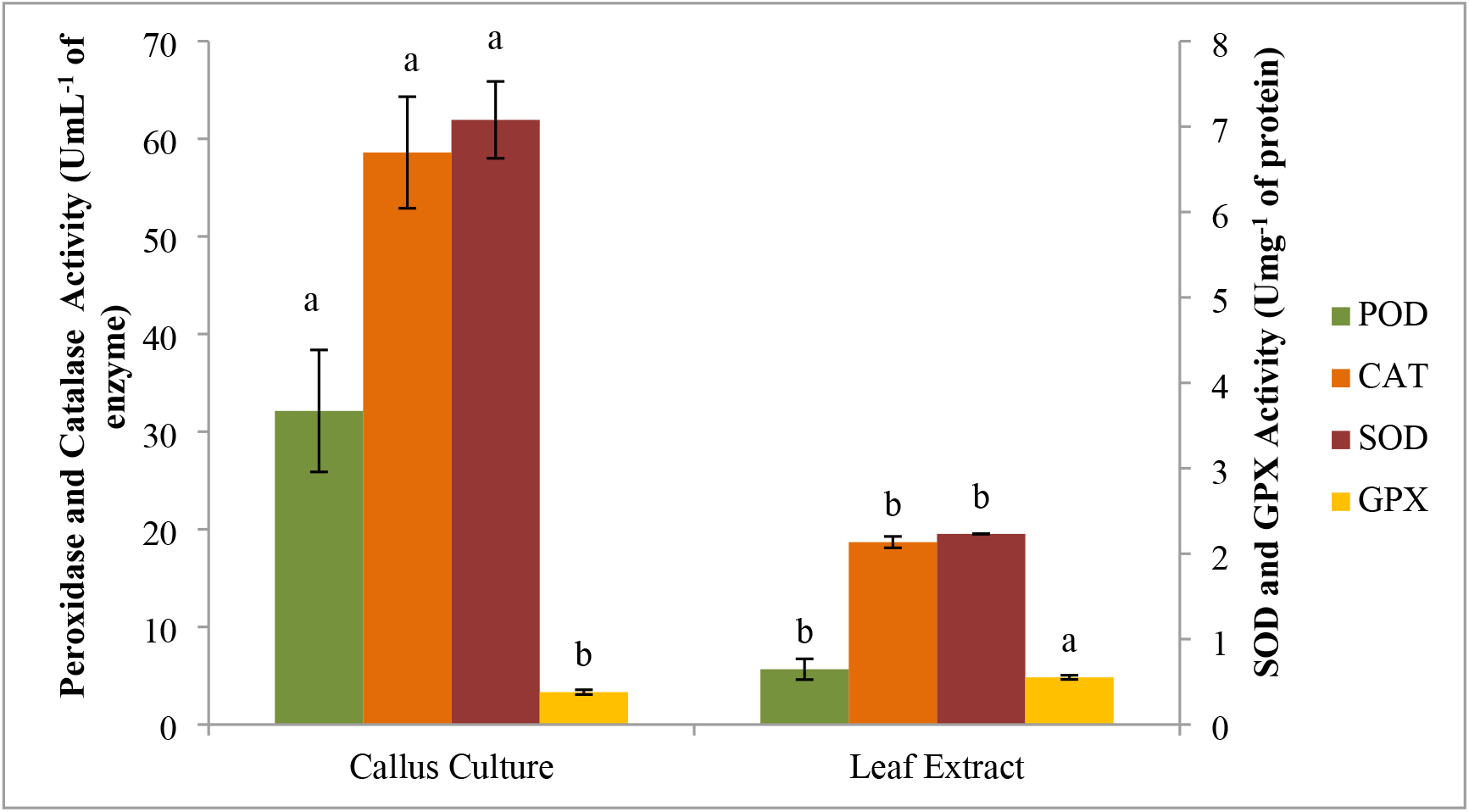
Comparative activity of antioxidants of callus culture and leaf extract of *C. camphora*.

**Figure 5:**
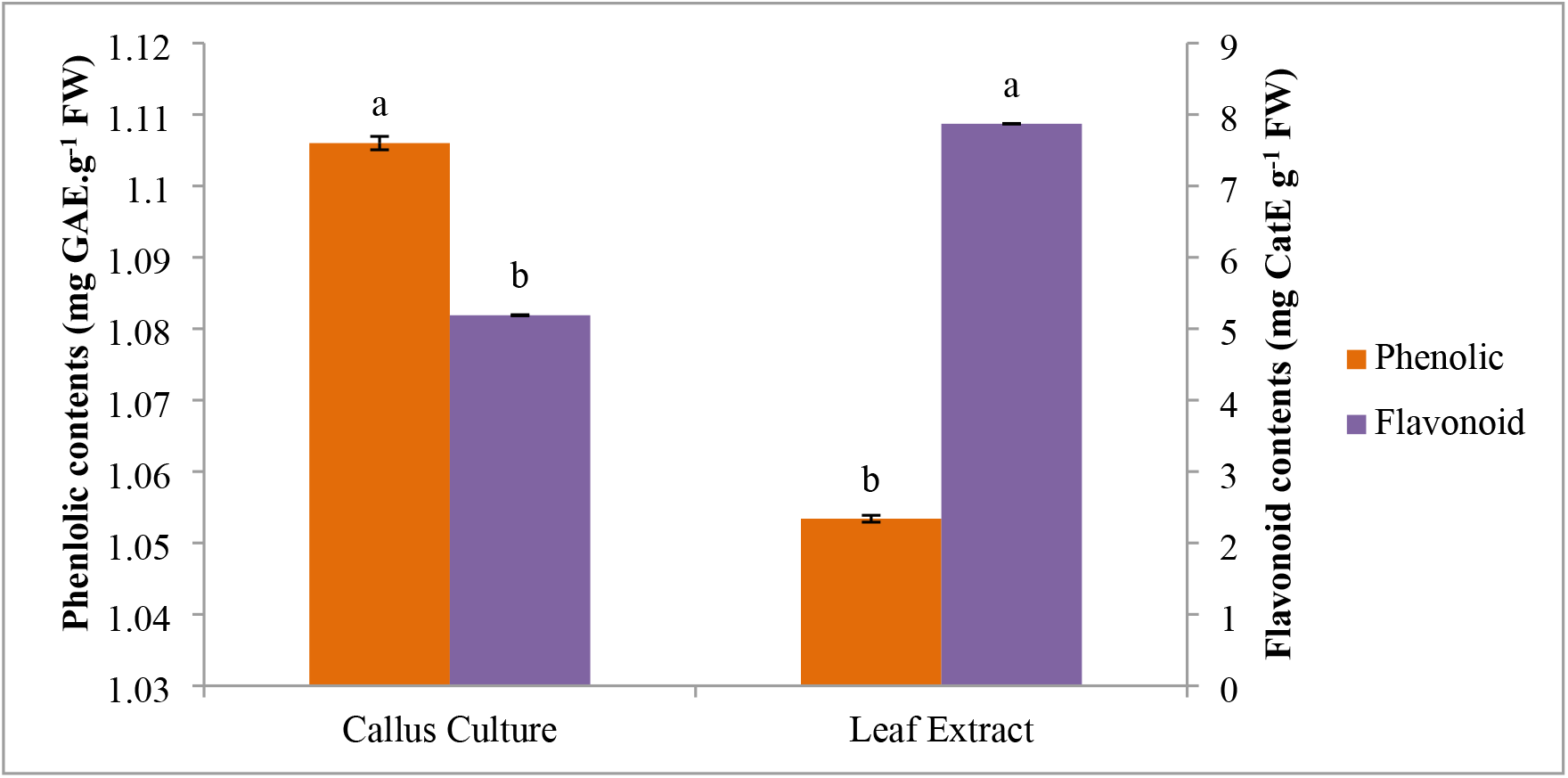
Comparative phenolic and flavonoid contents of callus culture and leaf extract *C. camphora*.

#### Total phenolic and flavonoid contents

The phenolic contents in callus culture were 1.106 mg GAE g^-1^ and in leaf extract were 1.053 mg GAE g^-1^. Overall, callus culture showed highest phenolic contents as compared to leaf extract. A decreasing trend in flavonoid contents was from 7.87 to 5.189 mg CatE g^-1^. Leaf extract of plant mentioned highest flavonoid contents (7.87 mg CatE g^-1^) while in callus cultures flavonoid contents were 5.189 mg CatE g^-1^ (Figure 6).

## DISCUSSION

In present study, two type of plant material (young leaves and nodal segments) were used as explant for shoot induction of camphor plant. Shoot initiation from nodal segments was achieved for establishment of *in vitro* plants. Different combinations of PGRs in MS media were tried for shoot induction. MS medium containing 2.26 µM 2, 4-D + 8.87 µM BAP provided best results with lateral shoot induction after 22 days as compared to other tested media. Similar results were recorded by Du *et al*. (2015) who observed that same combination and concentration of 2, 4-D and BAP for direct organogenesis by nodal explant from seedlings of cotyledonary embryo of camphor. It is worth mentioning here that 2, 4-D and BAP are more effective in triggering the cell elongation as compared to other growth regulators (Hemmati *et al*., 2020; Mayerni *et al*., 2020). When the combination of 0.4 µM TDZ and 2.26 µM 2, 4-D was used, shoots developed within 18 days without lateral shoot formation. Pakum *et al*. (2021) demonstrated similar results for direct organogenesis from leaf explant in *Kalanchoe tomentosa* Baker (Panda plant) on TDZ + 2, 4-D. Similarly, application of TDZ was also documented for camphor (Soulange *et al*., 2007). Lower level of TDZ (cytokinin) plays an important role in shoot induction and accelerates the plant regeneration (Kurup *et al*., 2018). BAP was previously used for shoot initiation of camphor explants (Haung *et al*., 1998; Babu *et al*., 2003; Sharma and Vashistha, 2010). During this investigation, MS medium with 4.92 µM IBA + AC 2.0 g L^-1^ was also showed 100% response of shoot initiation however, these results was contradictory to Babu *et al*. (2003), who reported rooting on this medium instead of shooting. The above-mentioned studies and our results confirmed that higher concentration of cytokinin’s with various concentrations of auxin supports the shoot induction.

In this investigation, the response of callus induction from young juvenile leaf and the nodal segments was also evaluated and found that significant and more positive response was observed on MS medium with combination of 8.87 µM BAP and 2.26 µM 2, 4-D as compared to other media. Similar results for callus initiation and proliferation from leaf disc of *C. camphora* were described by Chen *et al*. (2010) but in their case calli were induced on nodal explants instead of leaf segments. However, in another study, various combinations of auxins and cytokinin’s were describes for callus initiation from leaf explants of camphor (Soulange *et al*., 2007; Aref and Salem, 2020). Similarly, 2, 4-D and BAP was also reported as good for callus induction and proliferation in many other plant species. Since, combination of both synergistically more effective in promoting callus development by triggering the cell elongation (Hemmati *et al*., 2020; Mayerni *et al*., 2020; Bong *et al*., 2021; Lu *et al*., 2023; Muthi’ah *et al*., 2023). 2, 4-D alone was also suggested as the best auxin for callus induction and proliferation in many previous studies (Al-Ajlouni *et al*., 2015; Anjusha and Gangaprasad, 2017; Farhadi *et al*., 2017; Farvardin *et al*., 2017). In present study, the morphology of developed callus was lush green while compact in texture. Muthi’ah *et al*. (2023) observed similar morphology of callus on this combination of growth regulators in his study on *Calotropis gigantea*. Aref and Salem (2020) found different morphology of calli of camphor on different combinations of PGRs by using different explants. According to Ryu *et al*. (1992), even within the same genus, the appearance of callus depends upon the concentration and type of plant growth regulators. In present investigation, maximum proliferation response, growth index of callus (0.76g within 19 days) after callus sub-culturing was recorded on BAP + 2, 4-D in concentrations of 8.87 µM + 2.26 µM. Morphologically, callus was lush green, granular, lightly friable and compact in nature. El-Kader *et al*. (2019) reported similar results by sub-culturing of camphor leaf callus on MS medium fortified with 2.0 mg L^-1^ NAA and 1.0 mg L^-1^ BAP. However, contrary to our results, best yield of 11 g was obtained after 45 days of sub-culturing in their study. For high yield of callus various combinations of auxins and cytokinin’s were used in literature (Kintzios *et al*., 1999; Karam *et al*., 2003). Time interval of callus sub-culture varies among various species in literature also (Klimaszewska *et al*., 2016). These proliferated callus cultures were used for comparative analysis of phytochemical and antioxidant activity. Junairiah *et al*. (2019) and Astuti *et al*. (2020) reported that compact calli were far better regarding to production of secondary metabolites as compared friable ones.

In current study, a comparative analysis of phytochemicals and antioxidant activities was assayed in callus vs. leaf extract of camphor plant. Callus culture showed great potential to produce secondary metabolites *i*.*e*., phytochemicals and antioxidants those were also found in whole plants. Various investigations and approaches have been done for the production and to overcome the limitations of these vital secondary metabolites. For different applications of biotechnological fields these *in vitro* extract of callus cultures were utilized successfully (Fazal *et al*., 2016; Efferth, 2019; El-Kader *et al*., 2019; Aref and Saleem, 2020). In our findings, an increasing trend in the activities of peroxidase, catalase, and superoxide dismutase was observed in callus culture as compared to leaf extract. These results are in line with the previous study of Ali *et al*., (2021) who demonstrated that the application of GA_3_ (PGR) on *sorghum* seedling resulted in increasing SOD and POD activity. POD, SOD and catalase were considered as important enzymes which activate the defense mechanisms of plant by regulating metabolic processes (Liang *et al*., 2017; Huseynova *et al*., 2014). Reason behind this increasing trend might be the inhibition of reactive oxygen species (ROS) by a variety of phytochemicals and antioxidant enzymes (Namdeo, 2007), which are essential for protecting the calli cells from various stress factors. In present study, a decreasing trend was found in Glutathione peroxidase activity. In plant, GPXs may play a variety of functions in stress tolerance and growth (Sharma *et al*., 2021). It was reported that GPX activity results in accumulation of high rate of ROS in zygotic or embryonic nuclei (Rattanawong *et al*., 2021) and thought to have signaling factors during abiotic stress (Delaunay *et al*., 2002). This study was in line with Passaia *et al*. (2015) who described that GPXs control the plant growth and development in both favorable and unfavorable environments. SOD and GPx have the ability to directly balance the oxidative stress and provide protection to plant cells from DNA damage (Strycharz-Dudziak *et al*., 2020). SOD initially, catalases the dismutation of molecular oxygen (O_2_) and released hydrogen peroxide (H_2_O_2_) while the primary enzyme that breaks down this hydrogen peroxide into water in cells is glutathione peroxidase (Gurudath *et al*., 2012).

In present work, statistically significant increase in phenolic contents and decrease in flavonoid contents was recorded in callus cultures as compared to leaf extract grown in natural conditions. Medicinal plants with high concentrations of phenols and flavonoids have strong antioxidant effects. Our findings indicated that the phenylpropanoid pathway is activated by PGRs-containing media because this pathway causes an increase in synthesis of phenolic contents (Khan *et al*., 2016). It is reported that Phenylalanine ammonia-lyase (PAL) enzyme activity increased by application of plant growth regulators exogenously and it causes the accumulation of phenols and other secondary metabolites (Nagai *et al*., 1994; Khan *et al*., 2016). Therefore, present study revealed that phenylpropanoid molecule production was positively impacted by various concentrations of PGPRs and phenolic contents were produced in response of defense mechanism. Correlation between total phenolic content and antioxidants in other plants has been recorded (Pérez-Tortosa *et al*., 2012). According to Hatami *et al*. (2016), plant metabolism is quite complicated and depends on variety of variables, including the types and concentrations of growth hormones in the culture media, age, and type of cells or tissues. Overall results demonstrate that an increase in antioxidant activity and phenolic contents was due to various factors that might be the oxidative stress on *in vitro* callus cultures.

## CONCLUSION

In conclusion, it was observed during this investigation that callus cultures extract contains higher phytochemicals and antioxidant enzymes as compared to leaf extract of camphor plant and this enhanced activity of enzymatic antioxidants and phenolic contents might be correlated with *in vitro* conditions and PGRs used during callus induction. The results hint the potential use of callus cultures of *C. camphora* for the production of phytochemicals and antioxidants enzymes at commercial scale. These plant-based antioxidants and phytochemicals can be used in pharmaceutical, cosmetics and food industries to meet the country demand. However, this study requires more investigations to assess the efficiency of various combinations of plant growth regulators or some other elicitors to enhance these phytochemicals and enzymes under *in vitro* conditions.

## Availability of data and materials

All the data associated with this study are included in this manuscript.

## Authors contribution

**S.R** Perform the experiment and prepared the manuscript **SAK & ZAS** Supervise and proofread the manuscript.

## REFERENCES

Al-Ajlouni Z.I., Abbas S., Shatnawi M., Al-Makhadmeh I. (2015). In vitro propagation, callus induction, and evaluation of active compounds on Ruta graveolens. J Food Agricult Environ, 13(2):101–106.

Ali A. A., Ibrahim M. E. H., Li X., Nimir N. E. A., Elsiddig A. M. I., Jiao X., Zhu G., Salih E. G. I., Suliman M. E., Elradi S. B. M. (2021). Gibberellic acid and nitrogen efficiently protect early seedlings growth stage from salt stress damage in sorghum. Scie Rep, 11(1):66–72. 10.1038/s41598-021-84713-9

Anjusha S., Gangaprasad A. (2017). Callus culture and in vitro production of anthraquinone in Gynochthodes umbellata (L.) Razafim. & B. Bremer (Rubiaceae). Ind Crops Prod, 95:608– 614. 10.1016/j.indcrop.2016.11.021

Aref M. S., Salem S. S. (2020). Bio-callus synthesis of silver nanoparticles, characterization, and antibacterial activities via Cinnamomum camphora callus culture. Biocat Agricult Biotechnol, 27(2):101689. 10.1016/j.bcab.2020.101689

Aryal S., Baniya M. K., Danekhu K., Kunwar P., Gurung R., Koirala N. (2019). Total phenolic content, flavonoid content and antioxidant potential of wild vegetables from Western Nepal. Plants, 8(4):96. 10.3390/plants8040096

Astuti R. D., Harahap F., Edi S. (2020). Callus induction of mangosteen (Garcinia mangostana L.) in vitro with addition of growth regulators. Journal of Physics: Conference Series, 1485(1):012029. 10.1088/1742-6596/1485/1/012029

Babu K. S., Sajina A., Minoo D., John C. Z., Mini P., Tushar K. V., Rema J., Ravindran P. N. (2003). Micropropagation of camphor tree (Cinnamomum camphora). Plant Cell Tiss Organ Cult, 74(2):179–183. 10.1023/a:1023988110064

Beers R. F., Sizer I. W. (1952). A spectrophotometric method for measuring the breakdown of hydrogen peroxide by catalase. J Biol Chem, 195(1):133–140. 10.1016/s0021-9258(19)50881-x

Bhojwani S.S., Dantu P.K. (2013). Plant Tissue Culture: An Introductory Text. Tissue and cell culture. Springer eBooks, 39–50. 10.1007/978-81-322-1026-9

Bong F. J., Chear N. J., Ramanathan S., Mohana-Kumaran N., Subramaniam S., Chew B. L. (2021). The development of callus and cell suspension cultures of Sabah snake grass (Clinacanthus nutans) for the production of flavonoids and phenolics. Biocat Agricult Biotechnol, 33:101977. 10.1016/j.bcab.2021.101977

CABI (2022). ‘Cinnamomum camphora (camphor laurel)’, CABI Compendium. Centre for Agriculture and Bioscience International. doi: 10.1079/cabicompendium.13519

Chen J., Tang C., Zhang R., Ye S., Zhao Z., Huang Y., Xu X., Lan W., Yang D. (2020). Metabolomics analysis to evaluate the antibacterial activity of the essential oil from the leaves of Cinnamomum camphora (Linn.) Presl. J Ethnopharma, 253(1):112652. 10.1016/j.jep.2020.112652

Chen M., Ye Z., Ouyang S., Lin S., Shao A., Huang L. (2010). Callus induction of Cinnamonum camphora and formation of borneol. Zhongguo Zhong yao za zhi = Zhongguo zhongyao zazhi = China J Chinese Materia Medica, 35(5):558–560. 10.4268/cjcmm20100503

Chowdhary V., Alooparampil S., Pandya R. V., Tank J. G. (2022). Phenolic Compounds: Chemistry, Synthesis, Diversity, Non-Conventional Industrial, Pharmaceutical and Therapeutic Applications. Physiological function of phenolic compounds in plant defense system, edited by Farid A. Badria, Intech Open eBooks, 1, 1–22. 10.5772/intechopen.101131

Chun O. K., Kim D. (2004). Consideration on equivalent chemicals in total phenolic assay of chlorogenic acid-rich plums. Food Resh Inter, 37(4):337–342. 10.1016/j.foodres.2004.02.001

Cory H., Passarelli S., Szeto J., Tamez M., Champagne C. M. (2018). The Role of polyphenols in human health and food systems: A mini-review. Front Nutri, 5: 87. 10.3389/fnut.2018.00087

Delaunay A., Pflieger D., Barrault M., Vinh J., Toledano M. B. (2002). A thiol peroxidase is an H<sub>2</sub>O<sub>2</sub> receptor and redox-transducer in gene activation. Cell, 111(4):471–481. 10.1016/s0092-8674(02)01048-6

Donadio G., Mensitieri F., Santoro V., Parisi V., Bellone M. L., De Tommasi N., Izzo V., Piaz F. D. (2021). Interactions with microbial proteins driving the antibacterial activity of flavonoids. Pharmaceutics, 13(5):660. 10.3390/pharmaceutics13050660

Du L., Li Y., Yao Y., Zhang L. (2015). An efficient protocol for plantlet regeneration via direct organogenesis by using nodal segments from embryo-cultured seedlings of Cinnamomum camphora L. The Public Library of Science ONE, 10(5):1–10. 10.1371/journal.pone.0127215

Efferth T. (2019). Biotechnology applications of plant callus cultures. Engineering, 5(1):50–59. 10.1016/j.eng.2018.11.006

El-Kader E. M. A., Serag, A., Aref M. A., Ewais E. M., Farag M. A. (2019). Metabolomics reveals ionones upregulation in MeJA elicited Cinnamomum camphora (camphor tree) cell culture. Plant Cell Tiss Organ Cult, 137(2):309–318. 10.1007/s11240-019-01572-z

Farhadi N., Panahandeh J., Azar A. M., Salte S. (2017). Effects of explant type, growth regulators and light intensity on callus induction and plant regeneration in four ecotypes of Persian shallot (Allium hirtifolium). Sci Hort, 2(18):80–86. 10.1016/j.scienta.2016.11.056

Farvardin A., Ebrahimi A., Hosseinpour B., Khosrowshahli M. (2017). Effects of growth regulators on callus induction and secondary metabolite production in Cuminum cyminum. Nat Prod Res, 31(17):1963–1970. 10.1080/14786419.2016.1272105

Fazal H., Abbasi B. H., Ahmad N., Ali M. (2016). Elicitation of medicinally important antioxidant secondary metabolites with silver and gold nanoparticles in callus cultures of Prunella vulgaris L. Appl Biochem Biotechnol, 180(6):1076–1092. 10.1007/s12010-016-2153-1

Flohé L., Günzler W. A. (1984). Assays of glutathione peroxidase. Methods enzym, 105:114–121. 10.1016/S0076-6879(84)05015-1

Forni C., Facchiano F., Bartoli M., Pieretti S., Facchiano A., D Arcangelo D., Norelli S., Valle G., Nisini R., Beninati S., Tabolacci C., Jadeja R. N. (2019). Beneficial role of phytochemicals on oxidative stress and age-related diseases. BioMed Res Inter, 8748253. 10.1155/2019/8748253

Forni C., Frattarelli A., Lentini A., Beninati S., Lucioli S., Caboni E. (2016). Assessment of the antiproliferative activity on murine melanoma cells of extracts from elicited cell suspensions of strawberry, strawberry tree, blackberry and red raspberry. Plant Biosys, 150(6):1233–1239. 10.1080/11263504.2015.1018981

Gurudath S., Naik R., Ganapathy K., Guruprasad Y., Sujatha D., Pai A. (2012). Superoxide dismutase and glutathione peroxidase in oral submucous fibrosis, oral leukoplakia, and oral cancer: A comparative study. J Orofacial Sci, 4(2):114. 10.4103/0975-8844.106202

Hatami M., Kariman K., Ghorbanpour M. (2016). Engineered nanomaterial-mediated changes in the metabolism of terrestrial plants. Sci Total Environ, 571: 275– 291.

Hemmati, N., Cheniany, M., & Balestrini, R. (2020). Effect of plant growth regulators and explants on callus induction and study of antioxidant potentials and phenolic metabolites in Salvia tebesana Bunge. Botanica Serbica, 44(2):163–173. 10.2298/botserb2002163h

Huang L. C., Huang B. L., Murashige T. (1998). A micropropagation Protocol for Cinnamomum camphora. In Vitro Cell Develop Biol-Plant, 34(2):141–146.

Huseynova, I. M., Aliyeva, D. R., & Aliyev, J. A. (2014). Subcellular localization and responses of superoxide dismutase isoforms in local wheat varieties subjected to continuous soil drought. Plant Physiol Biochem, 81:54–60. 10.1016/j.plaphy.2014.01.018

Ivanova V., Stefova M., Vojnoski B., Dörnyei Á., Márk L., Dimovska V., Stafilov T., Kilár F. (2011). Identification of polyphenolic compounds in red and white grape varieties grown in R. Macedonia and changes of their content during ripening. Food Res Inter, 44(9):2851–2860. 10.1016/j.foodres.2011.06.046

Junairiah-Mahmuda A., Manuhara Y. S. W., Ni’matuzahroh., Sulistyorini L. (2019). Callus induction and bioactive compounds from Piper betle L. var nigra. Institute of Physics Publishing Conference Series: Earth Environ Sci, 217:1–9. 10.1088/1755-1315/217/1/012026

Kang N., Han S., Yoon S., Sim J., Maeng Y. H., Kang H., Yoo E. S. (2019). Cinnamomum camphora Leaves alleviate allergic skin inflammatory responses In Vitro and In Vivo. Toxicol Res, 35(3):279–285. 10.5487/tr.2019.35.3.279

Karam N., Jawad F., Arikat N., Shibli R. (2003). Growth and rosmarinic acid accumulation in callus, cell suspension, and root cultures of wild Salvia fruticosa. Plant Cell, Tiss Organ Cult, 73:117– 121.

Khan T. M., Abbasi B. H., Khan M. A., Shinwari Z. K. (2016). Differential effects of thidiazuron on production of anticancer phenolic compounds in callus cultures of Fagonia indica. Appl Biochem Biotechnol, 179(1):46–58. 10.1007/s12010-016-1978-y

Kintzios S., Nikolaou A., Skoula M. (1999). Somatic embryogenesis and in vitro rosmarinic acid accumulation in Salvia officinalis and S. fruticosa leaf callus cultures. Plant Cell Rep, 18(6):462–466. 10.1007/s002990050604

Klimaszewska K., Hargreaves C., Lelu-Walter M., Trontin J. (2016). Advances in conifer somatic embryogenesis since year 2000. In Methods in Molecular Biology, 131–166. 10.1007/978-1-4939-3061-6_7

Kurup S. S., Purayil F. T., Alkhaili M., Tawfik N. H., Cheruth A. J., Kabshawi M., Subramaniam S. (2018). Thidiazuron (TDZ) induced organogenesis and clonal fidelity studies in Haloxylon persicum (Bunge ex Boiss & Buhse): an endangered desert tree species. Physiol Mol Biol Plants, 24(4):683–692. 10.1007/s12298-018-0532-5

Lee H. S., Hyun E., Yoon W. J., Kim B., Rhee M. H., Kang H. G., Cho J. Y., Yoo E. S. (2006). In vitro anti-inflammatory and anti-oxidative effects of Cinnamomum camphora extracts. J Ethnopharma, 103(2):208–216. 10.1016/j.jep.2005.08.009

Lee O. N., Ak G., Zengin G., Cziáky Z., Jekő J., Rengasamy K. R., Park H. W., Kim D. H., Sivanesan, I. (2020). Phytochemical composition, antioxidant capacity, and enzyme inhibitory activity in callus, somaclonal variant, and normal green shoot tissues of Catharanthus roseus (L) G. Don. Molecules, 25(21):45–54. 10.3390/molecules25214945

Liang X., Chen Q., Lu H., Wu C., Lu F., Tang J. (2017). Increased activities of peroxidase and polyphenol oxidase enhance cassava resistance to Tetranychus urticae. Experi Appl Acarol, 71(3):195–209. 10.1007/s10493-017-0125-y

Lu X., Fei L., Li Y., Du J., Ma W., Huang H., Wang J. (2023). Effect of different plant growth regulators on callus and adventitious shoots induction, polysaccharides accumulation and antioxidant activity of Rhodiola dumulosa. Chinese Herbal Medi, 15(2):271–277. 10.1016/j.chmed.2022.07.005

Lucioli S., Di Bari C., Nota P., Frattarelli A., Forni C., Caboni E. (2017). Methyl jasmonate promotes anthocyanins’ production in Prunus salicina × Prunus persica in vitro shoot cultures. Plant Biosys, 151(5):788–791. 10.1080/11263504.2016.1255267

Luck H. (1974) Catalase. In Methods of Enzymatic Analysis, vol. II, edited by J. Bergmeyer & M, Grabi, Academic Press; New York, 885–890.

Luo Q., Xu C., Zheng T., Ma Y., Li Y., Zuo Z. (2021). Leaf morphological and photosynthetic differences among four chemotypes of Cinnamomum camphora in different seasons. Ind Crops Prod, 169:113651. 10.1016/j.indcrop.2021.113651

Mahajan S., Tuteja N. (2005). Cold, salinity and drought stresses: An overview. Arch Biochem Biophyss, 444(2):139–158. 10.1016/j.abb.2005.10.018

Maral J., Puget K., Michelson A. M. (1977). Comparative study of superoxide dismutase, catalase and glutathione peroxidase levels in erythrocytes of different animals. Biochem Biophys Res Commu, 77(4):1525–1535. 10.1016/s0006-291x(77)80151-4

Mayerni R., Satria B., Wardhani D., Chan S. (2020). Effect of auxin (2,4-D) and cytokinin (BAP) in callus induction of local patchouli plants (Pogostemon cablin Benth.). Institute of Physics Publishing Conference Series, 583(1):12–23. 10.1088/1755-1315/583/1/012003

McCown B.H., Lloyd G. (1981). Woody Plant Medium (WPM)—A mineral nutrient formulation for microculture of woody plant species. Hort Sci, 16: 453–453.

Muhamad S. H. A., On S., Sanusi S. N. A., Hashim A., Zai M. H. A. (2019). Antioxidant activity of Camphor leaves extract based on variation solvent. Journal of Physics: Conference Series, 1349(1):012102. 10.1088/1742-6596/1349/1/012102

Murashige T., Skoog F. (1962). A revised medium for rapid growth and bio assays with tobacco tissue cultures. J Plant Physiol, 15(3):473–497.

Muthi’ah A., Sakya A. T., Setyawati A., Samanhudi N., Rahayu M. (2023). Callus induction of Calotropis gigantea using BAP and 2,4-D in vitro. Institute of Physics Publishing: Conference Series, 1177(1):012021. 10.1088/1755-1315/1177/1/012021

Nagai N., Kitauchi F., Okamoto K., Kanda T., Shimosaka M., Okazaki M. (1994). A transient increase of phenylalanine ammonia-lyase transcript in kinetin-treated tobacco callus. Biosci, Biotechnol Biochem, 58(3):558–559. 10.1271/bbb.58.558

Namdeo A. (2007). Plant cell elicitation for production of secondary metabolites: A Review. Pharm Rev, 1:69–79

Nandi S., Vračko M., Bagchi M. C. (2007). Anticancer activity of selected phenolic compounds: Qsar studies using ridge regression and neural networks. Chem Biol Drug Des, 70(5):424–436. 10.1111/j.1747-0285.2007.00575.x

Ni Z., Han X., Chen C., Zhong Y., Xu M., Xu L., Yu F. (2021). Integrating GC-MS and ssRNA-Seq analysis to identify long non-coding RNAs related to terpenoid biosynthesis in Cinnamomum camphora. Ind Crops Prod, 171:113875. 10.1016/j.indcrop.2021.113875

Pakum W., Inmano O., Kongbangkerd A. (2021). TDZ and 2,4-D on in vitro propagation of panda plant from leaf explants. Orna Horticult, 27(1), 41–48. 10.1590/2447-536x.v27i1.2251

Passaia G., Margis-Pinheiro M. (2015). Glutathione peroxidases as redox sensor proteins in plant cells. Plant Sci, 234:22–26. 10.1016/j.plantsci.2015.01.017

Pérez-Tortosa V., López-Orenes A., Martínez-Pérez A., Ferrer M. T. E., Calderón A. (2012). Antioxidant activity and rosmarinic acid changes in salicylic acid-treated Thymus membranaceus shoots. Food Chem, 130(2):362–369. 10.1016/j.foodchem.2011.07.051

Ramakrishna A., Ravishankar G. A. (2011). Influence of abiotic stress signals on secondary metabolites in plants. Plant Signal Behav, 6(11):1720–1731. 10.4161/psb.6.11.17613

Rattanawong K., Koiso N., Toda E., Kinoshita A., Tanaka M., Tsuji H., Okamoto T. (2021). Regulatory functions of ROS dynamics via glutathione metabolism and glutathione peroxidase activity in developing rice zygote. Plant J, 108(4):1097–1115. 10.1111/tpj.15497

Roy A., Khan A., Ahmad I., Alghamdi S., Rajab B. S., Babalghith A. O., Alshahrani M. Y., Islam S., Islam M. R. (2022). Flavonoids a bioactive compound from medicinal plants and its therapeutic applications. BioMed Res Inter, 2022:1–9. 10.1155/2022/5445291

Ryu J. H., Doo H. S., Kwon T. H. (1992). Induction of haploid plants by another culture in sesame (Sesamum indicum L.) -(1)-effects of growth regulators and difference between genotypes on callus induction. Korean J Plant Tiss Cult (Korea Republic), 19: 171–177.

Samaniego I., Espin S., Cuesta X., Arias V., Rubio A., Llerena W., Angós I., Carrillo W. (2020). Analysis of environmental conditions effect in the phytochemical composition of potato (Solanum tuberosum) Cultivars. Plants, 9(7), 815. 10.3390/plants9070815

Shahidi F., Ambigaipalan P. (2015). Phenolics and polyphenolics in foods, beverages and spices: Antioxidant activity and health effects – A review. J Fun Foods, 18:820–897. 10.1016/j.jff.2015.06.018

Sharma H. (2021). In vitro propagation using nodal explants of Cinnamomum camphora: an important medicinal tree. Inter J Res Biosci Agricult Technol, 17: 394–401.

Sharma H., Vashistha B.D. (2010). In vitro propagation of Cinnamomum camphora (L.) nees & eberm using shoot tip explants. Ann Biol, 26(2): 109–114.

Soulange J.G., Sanmukhiya V.M.R. Seeburrm S.D. (2007). Tissue culture and RAPD analysis of Cinnamomum camphora and Cinnamomum verum. Biotechnology, 6: 239–244.

Strycharz-Dudziak M., Fołtyn S., Dworzański J., Kiełczykowska M., Malm M., Drop B. Polz-Dacewicz M. (2020). Glutathione peroxidase (GPx) and superoxide dismutase (SOD) in oropharyngeal cancer associated with EBV and HPV coinfection. Viruses, 12(9):1008. 10.3390/v12091008

Tajammal A., Siddiqa A., Al-Sehemi A. G., Azam M. A., Hafeez H., Munawar M. A., Basra M. A. R. (2021). Antioxidant, molecular docking and computational investigation of new flavonoids. J Mol Stru, 1254:132189. 10.1016/j.molstruc.2021.132189

Tian Z., Luo Q., Li Y., Zuo Z. (2020). Terpinene and β-pinene acting as signaling molecules to improve Cinnamomum camphora thermotolerance. Ind Crops Prod, 154, 112641. 10.1016/j.indcrop.2020.112641

Vignesh A., Subramaniam S., Vasanth K. (2022). Comparative LC-MS analysis of bioactive compounds, antioxidants and antibacterial activity from leaf and callus extracts of Saraca asoca. Phytomed Plus, 2(1):100167. 10.1016/j.phyplu.2021.100167

Wang J., Su B., Jiang H., Cui N., Yu Z., Yang Y., Sun Y. (2020a). Traditional uses, phytochemistry and pharmacological activities of the genus Cinnamomum (Lauraceae): A review. Fitoterapia, 146:104675. 10.1016/j.fitote.2020.104675

Wang L., Zhang K., Zhang K., Zhang J., Fu J., Li J., Wang G., Qiu Z., Wang X., Li J. (2020b). Antibacterial activity of Cinnamomum camphora essential oil on Escherichia coli during planktonic growth and biofilm formation. Front Microbiol, 11:561002. 10.3389/fmicb.2020.561002

Wei C., Li H., Cui G., Ma C., Deng R., Zou Z., Liu Z. (2022). Efficient separation of Cinnamomum camphora leaf essential oil and in vitro evaluation of its antifungal activity. Arab J Chem, 15(11):105225. 10.1016/j.arabjc.2022.104225

Zhang G., Yan X., Wu S., Ma M., Yu P., Gong D., Deng S., Deng S. (2020). Ethanol extracts from Cinnamomum camphora seed kernel: Potential bioactivities as affected by alkaline hydrolysis and simulated gastrointestinal digestion. Food Res Inter, 137:109363. 10.1016/j.foodres.2020.109363

